# Orthology clusters from gene trees with *Possvm*

**DOI:** 10.1101/2021.05.03.442399

**Authors:** Xavier Grau-Bové, Arnau Sebé-Pedrós

## Abstract

*Possvm* (<u>P</u>hylogenetic <u>O</u>rtholog <u>S</u>orting with <u>S</u>pecies o<u>V</u>erlap and <u>M</u>CL) is a tool that automates the process of classifying clusters of orthologous genes from precomputed phylogenetic trees. It identifies orthology relationships between genes using the species overlap algorithm to infer taxonomic information from the gene tree topology, and then uses the Markov Clustering Algorithm (*MCL*) to identify orthology clusters and provide annotated gene family classifications. Our benchmarking shows that this approach, when provided with accurate phylogenies, is able to identify manually curated orthogroups with high precision and recall. Overall, *Possvm* automates the routine process of gene tree inspection and annotation in a highly interpretable manner, and provides reusable outputs that can be used to obtain phylogeny-informed gene annotations and inform comparative genomics and gene family evolution analyses.

## Introduction

Gene orthology inference is a central problem in genomics and comparative biology (Koonin 2005). Orthology information can serve as the basis for gene family classification, make inferences about gene function under the ‘ortholog conjecture’ (Koonin 2005), enable cross-species comparisons, or trace the evolutionary dynamics of gene families, e.g. looking for specific expansions or secondary losses (Glover et al. 2019). In addition to genome-scale methods (Altenhoff et al. 2016; Glover et al. 2019), a common orthology inference strategy involves the supervised construction of phylogenies, followed by manual curation in order to make informed inferences about gene family evolution. Yet, supervised tree inspection can be labour-intensive and difficult to scale.

Here we present *Possvm* (<u>P</u>hylogenetic <u>O</u>rtholog <u>S</u>orting with <u>S</u>pecies o<u>V</u>erlap and <u>M</u>CL), a flexible and accurate tool to identify pairs and clusters of orthologous genes (orthogroups) from pre-computed phylogenies. It relies on the species overlap algorithm (Huerta-Cepas et al. 2007; Huerta-Cepas et al. 2016) to identify pairs of orthologous genes in a phylogeny, and processes this output to identify clusters of orthologs at various user-defined taxonomic depths, propagate gene name annotations, and report evolutionary relationships between gene pairs. *Possvm* can work with minimal input information: only a gene tree in *NEWICK* format, with or without node statistical supports. As the species overlap algorithm relies on the implicit taxonomic information contained in the gene tree’s topology, this approach is suitable for cases where the species tree is unknown or unavailable.

## New approaches

*Possvm* identifies pairs and clusters of orthologs (orthogroups) from a pre-computed gene tree, in three main steps (Figure 1A). First, *Possvm* takes as input a gene tree where species are specified as a prefix to the gene name (e.g. *species id. | gene id.*), and identifies pairs of orthologous genes using the species overlap algorithm (Huerta-Cepas et al. 2007; Huerta-Cepas et al. 2016). By default, no overlap in species composition is tolerated at any bipartition (species overlap threshold = 0), but this parameter can be adjusted (where greater overlap will result in more inclusive, less granular orthology calls). Second, *Possvm* builds a graph where pairs of genes (nodes) are linked according to their orthology relationships (edges). If tree bipartitions contain statistical supports (e.g. bootstraps or Bayesian posterior probabilities), this graph can be pruned to remove poorly-supported edges. This graph is partitioned into orthology clusters using the Markov Clustering Algorithm (*MCL*) (Enright et al. 2002) and a user-defined inflation parameter (default is *I =* 1.6). This clustering strategy is commonly applied to binary protein networks such as protein-protein interaction graphs (Vlasblom and Wodak 2009). Finally, the software outputs a table with pairs of orthologous genes (from step one), a table with the orthogroup membership for each gene and its statistical support (step three), and an annotated gene tree with orthogroup labels next to each gene.

**Figure 1.**
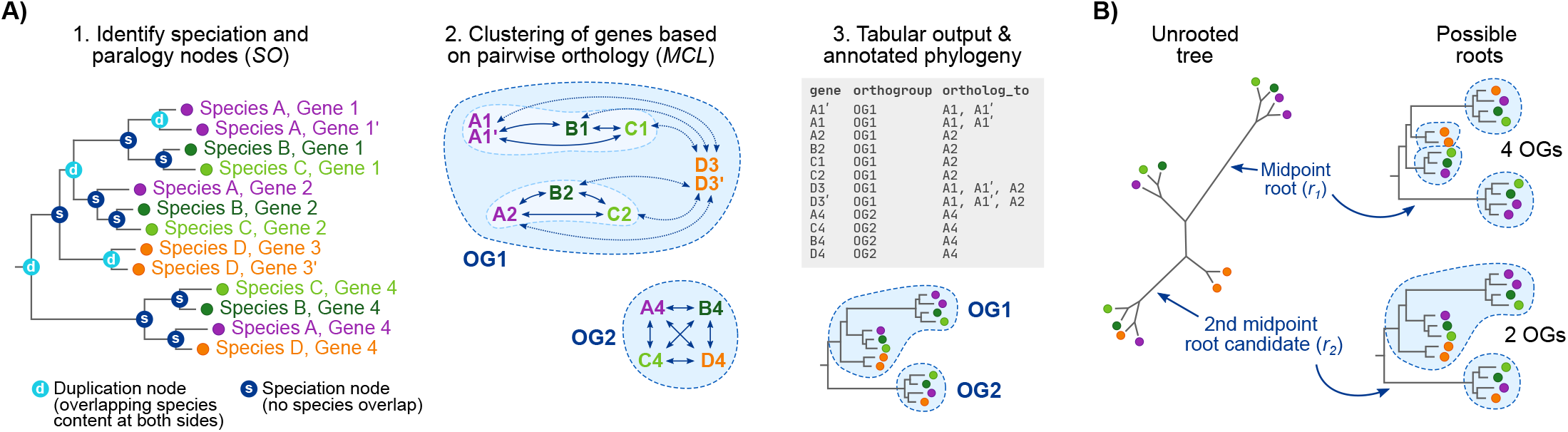
**A)** Summary of the main steps in *Possvm*. **B)** Example of the iterative mid-point rooting procedure. In this example, the original root (*r_1_*) results in the identification of four orthogroups whereas the second iteration (*r_2_*) results in two.

Optionally, the network clustering step (Figure 1A) can be tweaked to focus on a subset of species. This option allows the user to build gene trees with well-sampled taxonomic outgroups, while restricting the orthology-calling step to the ingroup of interest. Following the example in Figure 1A, the user could choose to include only species A, B and C, which would result in the split of OG1 into two smaller, more granular orthogroups (as orthology edges leading to species D are ignored). This principle can be applied systematically to obtain hierarchical orthogroups.

The species overlap algorithm requires a rooted gene tree in order to infer orthology relationships. Tree rooting can be done using *a priori* knowledge about taxonomic outgroups, using the mid-point rooting heuristics, or incorporating computationally intensive procedures such as non-reversible evolutionary models (Yap and Speed 2005; Bettisworth and Stamatakis 2020; Minh et al. 2020) and species tree reconciliation (Wu et al. 2014). *Possvm* offers the possibility to (i) use pre-computed rooted trees, (ii) perform mid-point rooting, or (iii) perform an iterative rooting procedure based on mid-point rooting, which selects a root based on an implicit parsimony criterion that minimises the number of ancestral gene copies in a given tree. In this iterative rooting approach (Figure 1B), *Possvm* will start by identifying the mid-point root (initial root, *r_1_*) and call orthogroups from that topology; then, it will ignore the initial root node and try the second best mid-point root candidate (*r_2_*), up to *n* times (*r_n_*). Finally, it will select the root node that minimises the number of orthogroups in the tree. The iterative rooting procedure can be suitable in cases where a long internal branch could be mistakenly selected as root by the standard mid-point approach.

In addition, *Possvm* can attempt to annotate genes and orthogroups using gene names from one or more reference species. For these steps, a dictionary file mapping the reference gene accession numbers (as used in the input gene tree) to the annotation of interest is required. First, individual genes are annotated as orthologous to one or more genes from the reference set. Second, the entire orthogroup is labelled according to the reference genes within, creating a composite name if it contains more than one reference gene (e.g. “*name A / name B*”). Finally, orthogroups lacking any reference gene can be annotated according to their closest labelled orthogroups according to the tree topology (receiving a label in the following format “*like: name A*”). If available, *Possvm* will also report the statistical support for the deepest node in the phylogeny that supports a given gene name annotation.

*Possvm* is freely available in Github (https://github.com/xgrau/possvm-orthology) under a GNU General Public License v3.0 license, together with test data and installation instructions. It requires Python 3 and the libraries *ETE*3 (Huerta-Cepas et al. 2016), *numpy* (Harris et al. 2020), *pandas* (McKinney 2010; Pandas development team 2021), *networkx* (Hagberg et al. 2008) and *markov_clustering* (Enright et al. 2002).

## Benchmarking the accuracy of the orthology clustering

We used *POSSVM* to classify orthologs in the ANTP homeobox class, a multi-gene family of transcription factors that is highly expanded in animals. This analysis allowed us to evaluate the accuracy of our orthology clusterings —using the manually curated ANTP families available in the HomeoDB database (Zhong and Holland 2011) as a reference—, and probe the evolutionary history of ANTPs.

Our analysis included whole-genome sequences from 14 bilaterians (including reference species such as *Homo sapiens* and *Drosophila melanogaster*), 12 cnidarians, and two placozoans (Supplementary Material S1). We used a standard pipeline commonly used in many gene family evolution studies: we used known ANTPs from the HomeoDB database to identify homologs in our genomes of interest using similarity searches (with *diamond* (Buchfink et al. 2014); 1,565 hits), built a multiple sequence alignment (*mafft* 7 *E-INS-i* (Katoh and Standley 2013)), and a maximum-likelihood phylogenetic tree (*IQ-TREE* 2 (Kalyaanamoorthy et al. 2017; Hoang et al. 2018; Minh et al. 2020); Supplementary Material S2).

We evaluated the accuracy of *POSSVM* against a curated classification of ANTP families in nine model bilaterian species, available in HomeoDB (six vertebrates and three insects). *POSSVM* classified the 45 ANTP families in this reference set with high precision (weighted mean = 0.98) and recall (weighted mean = 0.94; Figure 2A). Precision and recall values were comparable when evaluating the insect and vertebrate orthologs separately (Supplementary Material S3).

**Figure 2.**
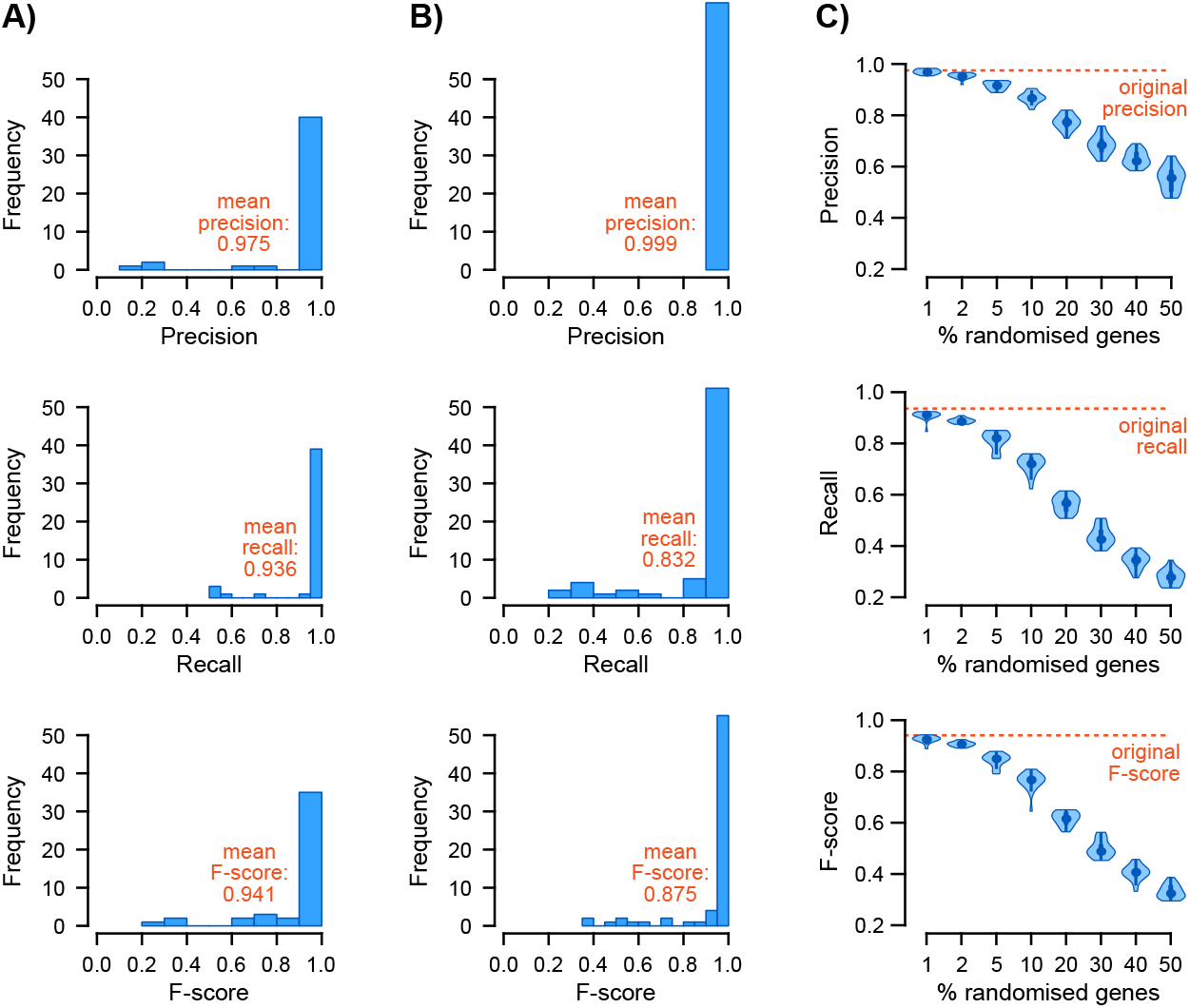
**A)** Precision, recall and F-score values for 45 ANTP families defined in HomeoDB. Mean values have been weighted by family size. **B)** *Id.* for 70 orthogroups defined in Orthobench. **C)** Effect of gene misplacement on precision, recall and F-score, for the ANTP dataset. Source data available in Supplementary Material S3.

Second, we evaluated *POSSVM*’s performance using 70 curated orthology groups from the Orthobench database, which were originally defined from manual examination of pre-computed gene trees (Trachana et al. 2011; Emms and Kelly 2020). In this dataset, *POSSVM* also exhibited very high precision across all reference orthogroups (weighted mean = 0.99), and comparable recall (weighted mean 0.83; Figure 2B and Supplementary Material S3).

Together, these results indicate that *Possvm* faithfully approximates the manual process of tree inspection aimed at orthogroup classification, identifying a single orthology group that perfectly matches the reference annotations in most cases (73% and 77% of the ANTP and Orthobench families, respectively).

Given *Possvm*’s reliance on pre-computed gene trees, its accuracy depends on the quality of the phylogenetic reconstruction. To evaluate how poorly constructed gene trees might affect *Possvm*’s orthology clusters, we randomised the position of tip nodes in the ANTP gene tree (Figure 2C). This analysis showed that, when we randomised up to 20% of node placements in the tree (i.e. 312 out of 1,565 genes), precision remained relatively high (above 0.75). On the other hand, gene tree inaccuracies have a strong detrimental effect on recall (*ca.* 0.5 at 20%).

## Phylogeny-informed gene annotation and evolutionary insights

*Possvm* is able to annotate orthogroups using gene names from a custom dictionary or a reference species. This functionality can be used to propagate gene annotations to non-model species in an orthology-aware manner, and inform the evolutionary history of a gene family. To illustrate this functionality, we annotated cnidarian ANTPs using human gene symbols as a reference (Figure 3). We find that *ca.* 50% of cnidarian ANTPs belong to orthogroups that can be labelled with one or more human genes within (Figure 3A, B). For example, out of 61 genes in the sea anemone *Nematostella vectensis*, *Possvm* annotates 33 genes as members of known ANTP families (Figure 3B). Among these, five are direct orthologs of a single reference gene (e.g. NKX3-2, Figure 3C), and 28 have one-to-many or many-to-many orthology relationships with human genes (e.g. *N. vectensis* encodes four co-orthologs of the human NKX2-2/NKX2-8 genes; Figure 3D).

**Figure 3.**
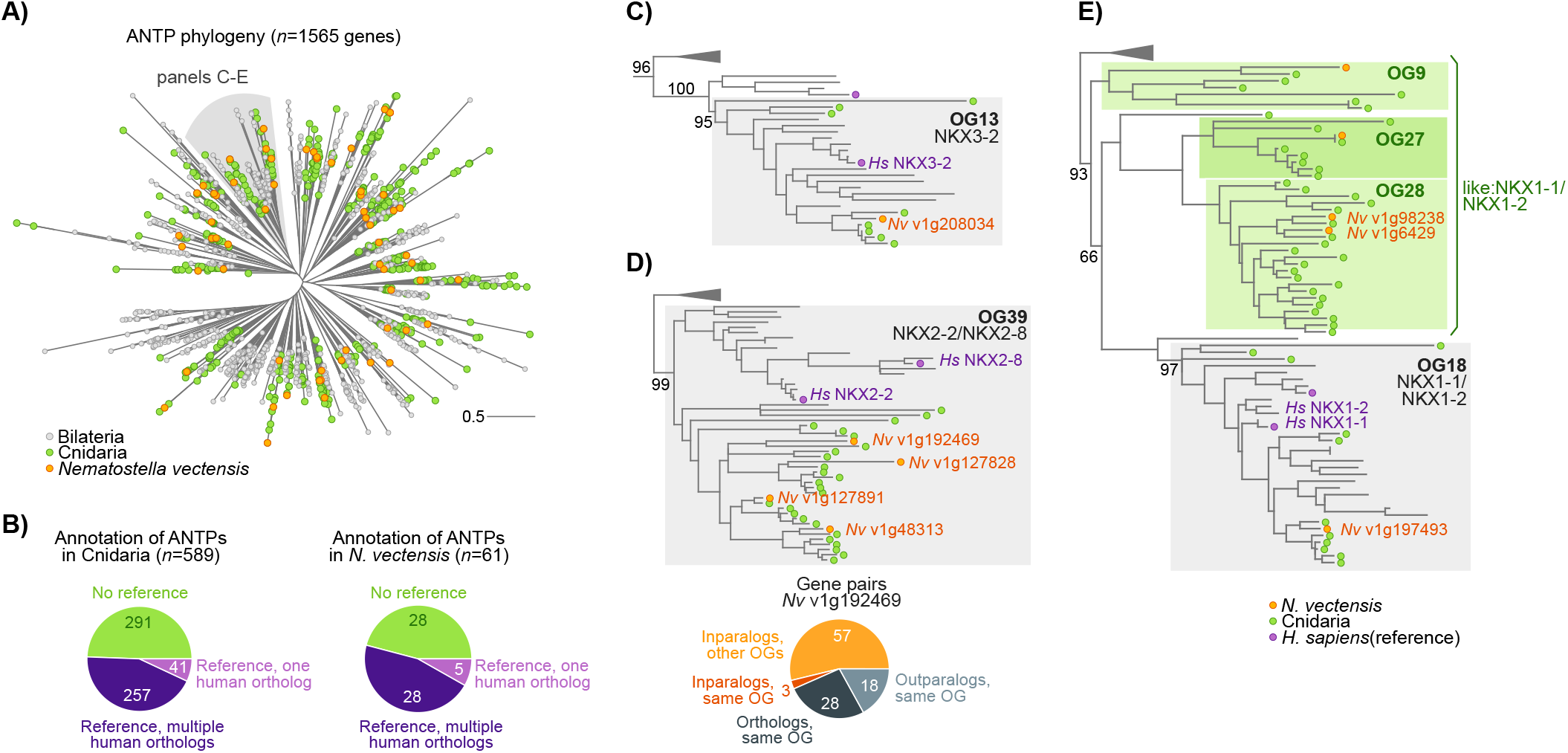
**A)** Global phylogeny of ANTP genes in bilaterians, cnidarians and placozoans. **B)** Summary of annotated genes in Cnidaria and in *N. vectensis*. **C-E)** Three examples of *Possvm* annotations from the ANTP phylogeny, including reporting evolutionary relationships at the gene pair level.

A further 28 *N. vectensis* ANTPs belong to orthogroups that could not be assigned to any known ANTP class based on their contents. Yet, *Possvm* is still able to label them as close paralogs of other orthogroups that do contain known genes, by propagating the annotations of phylogenetically close known orthogroups. Oftentimes, these unannotated orthogroups genes reflect cnidarian-specific duplicates with a many-to-one relationships with known ANTP families. For example, the NKX1-1/NKX1-2 family, which contains both bilaterian and cnidarian homologs (Figure 3E), is closely related to three cnidaria-specific orthogroups that would be annotated as similar to NKX1-1/NKX1-2 by *Possvm* (labelled as ‘*like: NKX1-1/NKX1-2*’). The greediness of this annotation propagation procedure can be controlled by defining a minimum statistical support for the last common ancestor between the annotated and unannotated orthogroups.

Finally, *Possvm* can also report fine-grained evolutionary relationships at the gene level. For example, taking as a reference the *N. vectensis* gene *v1g192469* (NKX2-2/NKX2-8 family), *Possvm* classifies its homologs as orthologs, in-paralogs or out-paralogs, within or without the same orthogroup (Figure 3D). By systematically reporting such relationships, we can dissect sets of homologous genes into precisely defined groups according to their evolutionary histories. This functionality allows the researcher to identify specific evolutionary patterns (e.g. intra-orthogroup duplications in a specific species), or to address evolutionary hypotheses in cross-species comparisons (e.g., testing the functional conservation of orthologous gene pairs compared to closely related paralogs).

## Discussion

*Possvm* is an accurate tool to automate the process of phylogeny parsing, ortholog clustering and gene name annotation propagation, requiring a gene tree as its sole input. Importantly, the species overlap algorithm (Huerta-Cepas et al. 2007; Huerta-Cepas et al. 2016) that sits at its core emulates an common heuristics used by researchers when inspecting a gene tree: it is assumed that a certain degree of taxonomic coherence should be present within an orthology group, but that small-scale inaccuracies in the tree inference might introduce discrepancies with the underlying species phylogeny. Therefore, these orthology classifications are highly interpretable when visualised over the gene tree —that can be produced by *Possvm*, together with table-based annotations—, which facilitates its critical appraisal by the researcher.

We have demonstrated that *Possvm* classifications show very high precision and high recall against a notably large multi-gene family (ANTP homeoboxes) and a curated benchmark of orthology groups (Trachana et al. 2011; Emms and Kelly 2020). Yet, it is crucial to highlight that *Possvm*’s performance depends on the quality of the input gene tree. In that regard, we have demonstrated that by combining the species overlap algorithm with *MCL* clustering we can tolerate relatively high rates of gene misplacement in the phylogenies and still maintain reasonable precision (Figure 2C).

In recent years, we have witnessed a rapid increase in the taxonomic sampling and quality of whole-genome sequencing efforts. Similarly, functional genomics data such as single-cell transcriptomic atlases are now available for diverse species (Plass et al. 2018; Sebé-Pedrós, Chomsky, et al. 2018; Sebé-Pedrós, Saudemont, et al. 2018; Musser et al. 2019). In that regard, *Possvm* provides an accurate and interpretable gene orthology inference solution that will facilitate gene family evolution studies, cross-species data integration, and large-scale comparative and functional genomics analyses.

## Supplementary material

**Supplementary Material S1.A)** Taxon sampling used in the ANTP phylogenetic analysis, with data sources. **B)** *Possvm* classification of metazoan ANTP genes.

**Supplementary Material S2.**Annotated maximum-likelihood phylogeny of metazoan ANTPs (*n =* 1565 genes), built with *IQ-TREE* (JTT+G4 model). Clades are color-coded according to *Possvm* annotations. Statistical supports obtained from 1,000 UF bootstrap iterations.

**Supplementary Material S3.A-D)** Detailed report of the precision, recall and F-scores of the ANTP (all 45 HomeoDB families, and the vertebrate- and insect-only analyses) and the Orthobench (70 families) reference datasets. For each orthology group defined in the original reference datasets, we indicate its size, the best *Possvm*-derived OG, and the precision, recall, and F-score statistics.

