## Supplementary figures and images for "Orthology clusters from gene trees with *Possvm*"

### Supplementary Material S2

**Supplementary Material S2**

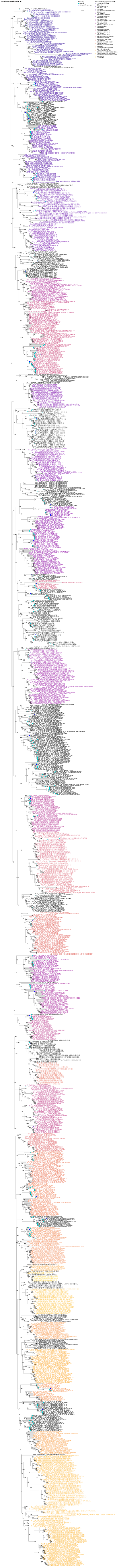
